# Rbec: a tool for analysis of amplicon sequencing data from synthetic microbial communities

**DOI:** 10.1101/2021.01.15.426834

**Authors:** Pengfan Zhang, Stjin Spaepen, Yang Bai, Stephane Hacquard, Ruben Garrido-Oter

## Abstract

**Summary:** Synthetic microbial communities (SynComs) constitute an emerging and powerful tool in biological, biomedical, and biotechnological research. Despite recent advances in algorithms for the analysis of culture-independent amplicon sequencing data from microbial communities, there is a lack of tools specifically designed for analysing SynCom data, where reference sequences for each strain are available. Here we present Rbec, a tool designed for the analysis of SynCom data that accurately corrects PCR and sequencing errors in amplicon sequences and identifies intra-strain polymorphic variation. Extensive evaluation using mock bacterial and fungal communities show that our tool outperforms current methods for samples of varying complexity, diversity, and sequencing depth. Furthermore, Rbec also allows accurate detection of contaminants in SynCom experiments.

**Availability and implementation:** Rbec is freely available as an open-source multi-platform R package. Release versions can be obtained via Bioconductor. The developer version is maintained and can be downloaded at: https://github.com/PengfanZhang/Rbec.

## 1 Introduction

Amplicon sequencing is a powerful technique to characterize microbial composition from environmental samples. Recent advances in algorithms and tools for the analysis of marker gene amplicon data have driven a shift from clustering approaches, based on operational taxonomic units (OTUs) and arbitrary sequence similarity thresholds, to error correction methods (Callahan *et al.*, 2016; Edgar and Flyvbjerg, 2015; Amir *et al.*, 2017; Peng and Dorman, 2020) that seek to estimate abundances of individual amplicon sequence variants (ASVs). A new generation of integrated pipelines (Bolyen *et al.*, 2019) allows researchers from a variety of fields in the environmental, biological, and medical sciences to reproducibly analyse marker gene sequencing data.

Synthetic microbial communities (SynComs) constitute an emerging and powerful tool to build experimentally tractable, reproducible microbial systems in the laboratory that enable controlled perturbation experiments and testing of falsifiable hypotheses. These reductionist approaches are being increasingly employed in studies of microbial ecology and evolution (Cairns *et al.*, 2020), plant and animal microbiota (Bai *et al.*, 2015; Zhang *et al.*, 2019; Vrancken *et al.*, 2019), and biotechnology (McCarty and Ledesma-Amaro, 2019). A factor limiting these new experimental approaches from developing to their full potential is the lack of bioinformatic tools specifically designed for the analysis of sequencing data obtained from gnotobiotic systems and SynComs. As a result, researchers typically employ standard clustering, error correcting or mapping approaches that do not take full advantage of these tractable experimental systems (e.g., reduced complexity and the availability of reference sequences for classification), resulting in reduced resolution, accuracy or data loss. To address this limitation, we developed a reference-based error correction algorithm, named Rbec, that is able to accurately and precisely correct PCR and sequencing errors in SynCom amplicon data, identify intra-strain polymorphism, and detect the presence of contaminants in gnotobiotic systems.

## 2 Methods

Rbec identifies and corrects sequencing errors by implementing a modified version of the quality-aware model implemented in the DADA2 tool (Callahan *et al.*, 2016). Rbec is specifically designed to efficiently and accurately process data from SynComs, for which reference sequences of individual community members are available (Fig. S1). First, reads are de-replicated into unique tags and subsequently aligned to the reference database, after which initial abundances are assigned to each strain according to the copy number of each exactly aligned tag. Next, tags that are not exactly matched to any sequence in the database are assigned a candidate error-producing reference based on *k*-mer distances. Sequencing reads are then subsampled and an error matrix is calculated using the mapping between subsampled reads and candidate error-producing sequences. The probability that a unique tag is erroneously produced by a given candidate error-producing sequence is calculated using a Poisson distribution. The probability and expectation values of this distribution are then used to determine whether a unique tag can be corrected from a reference sequence, and tags that cannot be corrected are subsequently removed. The parameters of the error model are recomputed iteratively until the number of re-assignments falls below a set threshold. Strain abundances are then estimated from the number of error-corrected reads mapped to each reference sequence. Finally, putatively contaminated samples are identified by a significant deviation from the expected proportion of corrected reads.

## 3 Results

To validate the performance of Rbec, we employed mock samples generated using a taxonomically wide set of 236 bacterial and 97 fungal strains obtained from sequenced culture collections derived from the *Arabidopsis thaliana* microbiota (Bai *et al.*, 2015; Durán *et al.*, 2018). Mapping of amplicon reads to the reference sequences showed that only 31.8% of all reads per sample, on average, had a perfect match in the database, indicating the presence of extensive sequencing and PCR errors and polymorphic copies (Fig. 1A, Fig. S2). Our implementation of the Rbec algorithm successfully corrected most erroneous reads (89.2% on average), out-performing all other tested *de novo* correction methods (Fig. 1B). This improvement was most pronounced for reads generated from polymorphic copies of the marker sequences within a single strain, owing to the fact that Rbec is capable of correctly classifying paralogous sequences (Fig. S3). To evaluate the accuracy of Rbec in characterizing community composition, we simulated *in silico* bacterial and fungal mock samples by mixing reads generated from sequencing individual isolates separately. For these simulations, we varied community complexity, strain similarity and sequencing depth. Across these three parameters, Rbec consistently performed better than all other tested methods in characterizing microbial composition in terms of deviation from the ground truth (Fig. 1C, S4 and S5), as well as precision and recall (Fig. S6), and was capable of detecting contaminated samples (Fig. S7).

**Figure 1.**
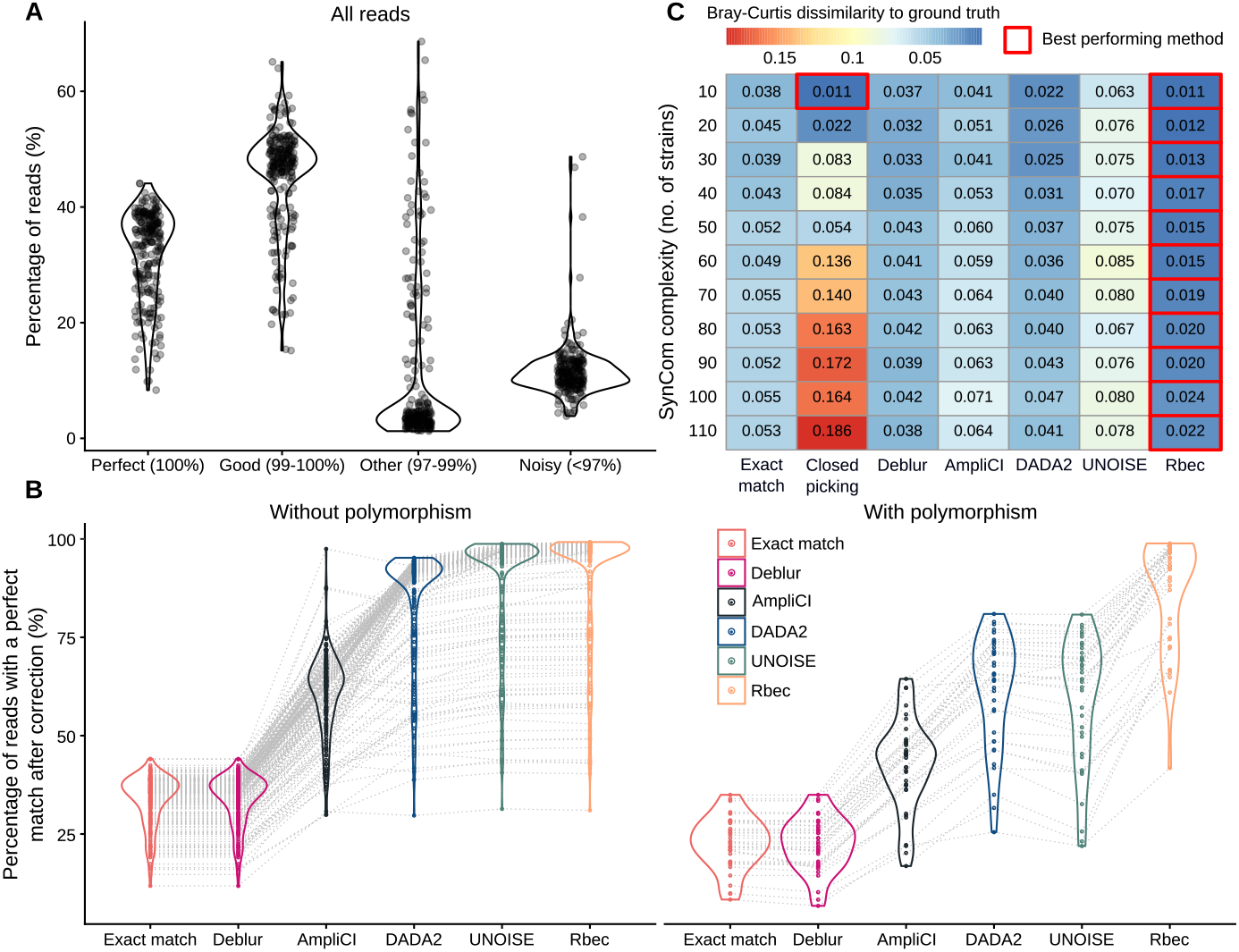
Evaluation of the Rbec algorithm. (**A**) Error profiles of amplicon sequencing data from 234 bacterial strains sequenced on the Illumina platform. (**B**) Percentage of perfectly aligned reads after correction with different methods. (**C**) Evaluation of the influence of community complexity on the performance of different methods, measured as a deviation from the ground truth using Bray-Curtis dissimilarities. The column names represent the number of strains in the mock communities and the row names represent the methods implemented to analyze the mock data. The values inside the heatmap refer to the averaged Bray-Curtis dissimilarities over 20 replicates for each threshold.

## Acknowledgements

We would like to thank Prof. Alga Zuccaro, Prof. Eric Kemen, and Dr. Yulong Niu for their feedback on this project.

