## Supplementary Materials for "Rbec: a tool for analysis of amplicon sequencing data from synthetic microbial communities"

**Appendix**

Details of Rbec algorithm

Library construction and preparation

Data processing with different methods

Simulations of mock communities

Detection of contamination sequences

Supplementary Figures

References

**Rbec algorithm**

A schematic workflow of Rbec is shown in Fig. S1. First, each reference sequence is assigned an initial abundance by counting the number of merged reads with an identical match. Then, merged reads in a sample are subsampled and subsequently *k*-mer distances between subsampled reads and each reference sequence are calculated. Next, the reference sequence showing the lowest *k*-mer distance to the query sequence is marked as the candidate error-producing sequence from which the query originates. If multiple candidates with the same *k*-mer distance are found, only the reference sequence with the highest initial abundance is considered as the original error-generating sequence. Next, the alignment results between the query reads and candidate error-producing references are used to derive the transition matrix. The transition matrix is a 20 by 43 matrix where the rows represent the transition combinations (e.g., A->A, A->T, A->G, A->C, T->T, …, -->G, -->C, including insertions), and the columns represent the sequence quality scores. Entries in the transition matrix are calculated by counting the number of each transition combination along the length of the alignment. The log-transformed transition matrix is then fitted with a weighted loess function to generate the error matrix.

Next, merged reads are dereplicated and the sequence quality scores for a unique tag are averaged over the scores of all identical copies of that sequence. Then, each unique tag is aligned to the reference database and the candidate error-producing sequence is identified as described above. We assume that the mismatches between query and reference are generated independently, so the rate at which a sequence *i* is produced from the error-generating reference *j* is calculated by the product over the error probabilities at the alignment positions where mismatches occur:

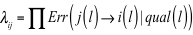

Similarly to the error-aware model implemented in DADA2 (Callahan *et al.*, 2016), the abundance probability of each unique tag is calculated using the *Poisson* distribution:

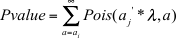

Where *E* is the expectation of the *Poisson* distribution, and *a'* is a cumulative abundance obtained by aggregating the abundances of unique tags assigned to a reference sequence. The tags with *P* values lower than 10^-40^, and an expectation lower than 0.05, are discarded. This expectation cut-off is intended to retain tags that could be produced at least once by the reference with the probability above 5%. The aim of this step is to retain tags that are generated from intra-strain amplicon sequence variants, which show high abundance relative to the reference sequence but do not exceed the *P* value cut-off. Reads above this threshold are then subsampled and aligned to the reference sequences in an iterative process. The iterations continue until the number of corrected reads falls below a fixed threshold, reaching convergence.

**Sequencing library construction and preparation**

To evaluate the errors in the output from sequencing and test this algorithm, we performed amplicon sequencing on 236 single bacterial strains and 97 fungal strains isolated from the root of *Arabidopsis thaliana* separately (Bai *et al.*, 2015; Durán *et al.*, 2018). Genomic DNA was isolated from each strain using the MP Biomedicals FastDNATM Spin Kit for Soil. DNA concentration was determined fluorometrically using the Quant-iT PicoGreen dsDNA Assay Kit (Thermo Fisher Scientific). The V5/V7 region of the 16S rRNA gene in bacteria and ITS1 region in fungi was amplified using the AACMGGATTAGATACCCKG (799F) and ACGTCATCCCCACCTTCC (1192R) primers, and CTTGGTCATTTAGAGGAAGTAA (ITS1F) and GCTGCGTTCTTCATCGATGC (ITS2R) primers, respectively, and indexing was done using Illumina-barcoded primers. The indexed amplicons were subsequently pooled, purified, and sequenced on the Illumina MiSeq platform.

To exclude the possibility that the observed error distribution of amplicon sequencing is specific to the MiSeq platform, we also analysed the amplicon sequencing data obtained using a HiSeq platform (Guo *et al.*, 2020).

**Data processing with different methods**

Raw reads were merged using Flash2 (-m 0.25 -M 250) (Magoc and Salzberg, 2011). Merged reads with ambiguous bases were excluded with USEARCH (Edgar, 2010). DADA2 and Deblur plug-ins in QIIME2 (Bolyen *et al.*, 2019) were applied to the filtered data by following the protocols indicated on the QIIME2 website (https://docs.qiime2.org/2019.7/tutorials/), to correct the reads and generate ASV tables. We also included the recently published algorithm AmpliCI (Peng and Dorman, 2020) for comparison purposes. We followed the instructions on the corresponding Github website (https://github.com/DormanLab/AmpliCI) to process the data and generate the ASV table. The abundances of ASVs showing exact matches to the reference sequences were extracted from this feature table. For the UNOISE tool (Edgar and Flyvbjerg, 2015), filtered reads were dereplicated by USEARCH and subsequently denoised using the -unoise3 function in USEARCH. For the exact match method, the filtered sequencing reads were aligned to the reference database by running the -uparse_ref command in USEARCH. Only hits with 100% identities were retained for generating the profiling table. The Deblur and AmpliCI methods were not applied to fungal data since the two methods require the equal length of input data, while the length of ITS sequences shows a large variation among strains.

**Simulation of mock communities**

To evaluate the performance of different algorithms when analysing with SynCom data with different complexities, strain similarities, and sequencing depths, we simulated mock samples using data obtained from sequencing clonal cultures individually. Firstly, the reference sequences of the V5-V7 region from all the strains were dereplicated, resulting in 114 strains having unique sequences in the V5-V7 region. To generate mock samples with different complexities, 10 to 110 strains from the candidate list containing 114 strains were randomly picked for each mock with a step size of 10 strains. The relative abundance of each strain was simulated using a log normal distribution (s.d. = 2). The total number reads in each mock was fixed at 10,000 reads and the reads for each strain were subsampled using Seqkit (Shen *et al.*, 2016) from the amplicon sequencing output of each individual strain, according to its abundance in the mock. To generate mock samples with different strain similarities, we set a maximum pairwise similarity threshold between each pair of strains in each mock community, ranging from 85% to 100%. To alleviate the influence of uneven abundance distribution of each strain on the evaluation, only 20 strains with equal abundances were included in mock communities. To evaluate the impact of different sequencing depths on the performance of the different algorithms, we simulated mock data with 50 strains at different depths, ranging from 500 to 10,000 reads. For each evaluation category, we generated 20 replicates for each parameter combination. 97 fungal strains were set as seeds to be randomly chosen to generate the fungal mock communities in the same way that we generated the bacterial mock communities. We applied the DADA2, UNOISE, Deblur, exact match, closed OTU picking, and Rbec methods to the simulated data sets. Finally, we calculated the Bray Curtis dissimilarities between the predicted profiling tables from different methods and the real composition for each mock sample (ground truth) using the vegan R package (Oksanen *et al.*, 2020).

**Detection of contamination sequences**

Existing error-correction algorithms cannot accurately estimate the abundances of strains with marker gene paralogs, and show a strong bias towards underestimation of their abundances. When analysing amplicon sequencing data obtained from synthetic communities, this leaves out numerous reads from paralog sequences, leading to low percentages of aligned reads per sample, which can be mistaken for the presence of contaminants. As Rbec not only corrects most erroneous amplicon sequencing reads, but also successfully identifies paralogous sequences, we expect that a high proportion of uncorrected reads is likely the result of contamination. Based on this assumption, Rbec includes a function to detect the contaminated samples and output the potential contamination sequences.

A sample *i* is flagged as contaminated, if

$$R_{i}<\mu-1.5*IQR$$

Where $R_{i}$ is the recruitment ratio of reads of sample *i*, $\mu$ is the mean of recruitment ratio of reads across samples, and $IQR$ is the interquartile range of the recruitment ratio. If a sequence accounts for more than 3% of total reads after error correction, we assume this sequence originates from a contaminant.

**Data deposition**

Raw sequencing data used to generate mock bacterial and fungal community data were deposited into the European Nucleotide Archive (ENA) under the accession number PRJEB43511. The scripts used for the computational analyses described in this study are available at https://github.com/PengfanZhang/Rbec, to ensure replicability and reproducibility of these results.

**Supplementary Figures**

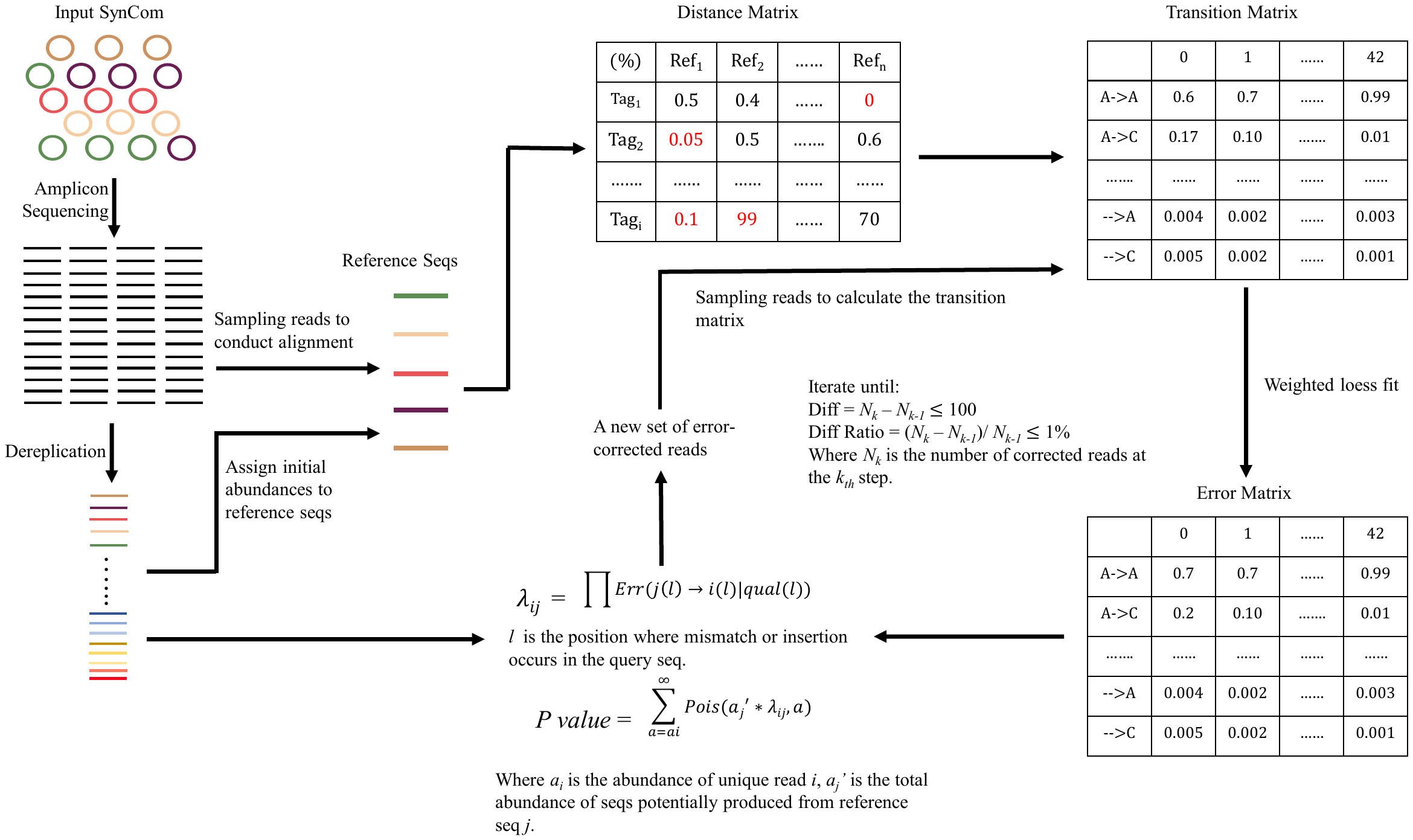

**Fig. S1. Schematic diagram of the Rbec algorithm.** Rbec consists of two main steps: error matrix estimation and abundance probability calculation. For the error matrix estimation, Rbec traverses through all query reads and reference sequences and matches each read with a unique candidate error-producing reference. Alignments between input and reference sequences are then used to calculate the error matrix. Abundance probabilities are then estimated by fitting a *Poisson* distribution.

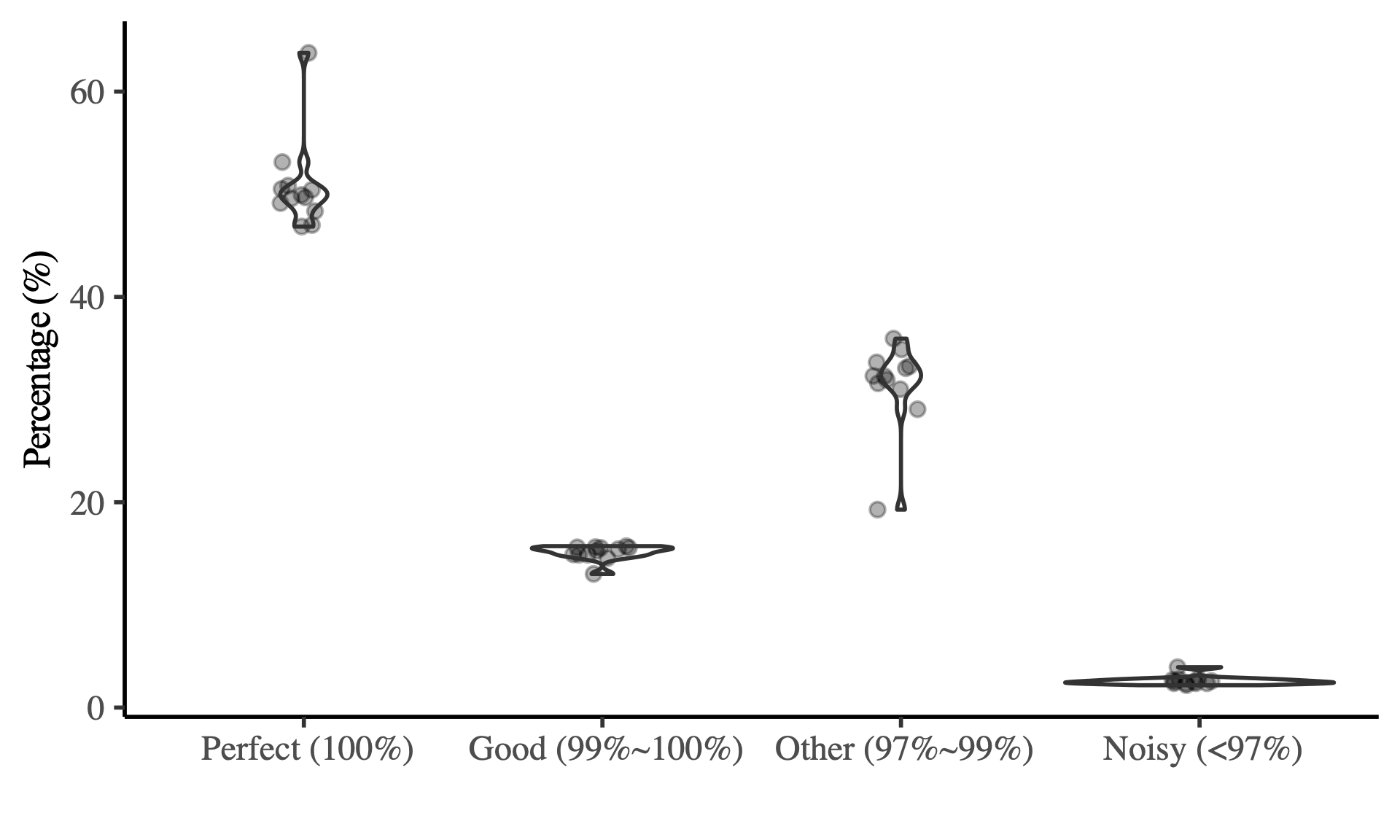

**Fig S2. Error distribution of amplicon sequencing data from a HiSeq sequencer.** Error profiles of amplicon sequencing data from 12 SynCom samples comprised of 12 bacterial strains and an artificial spike-in plasmid sequenced using the Illumina HiSeq platform.

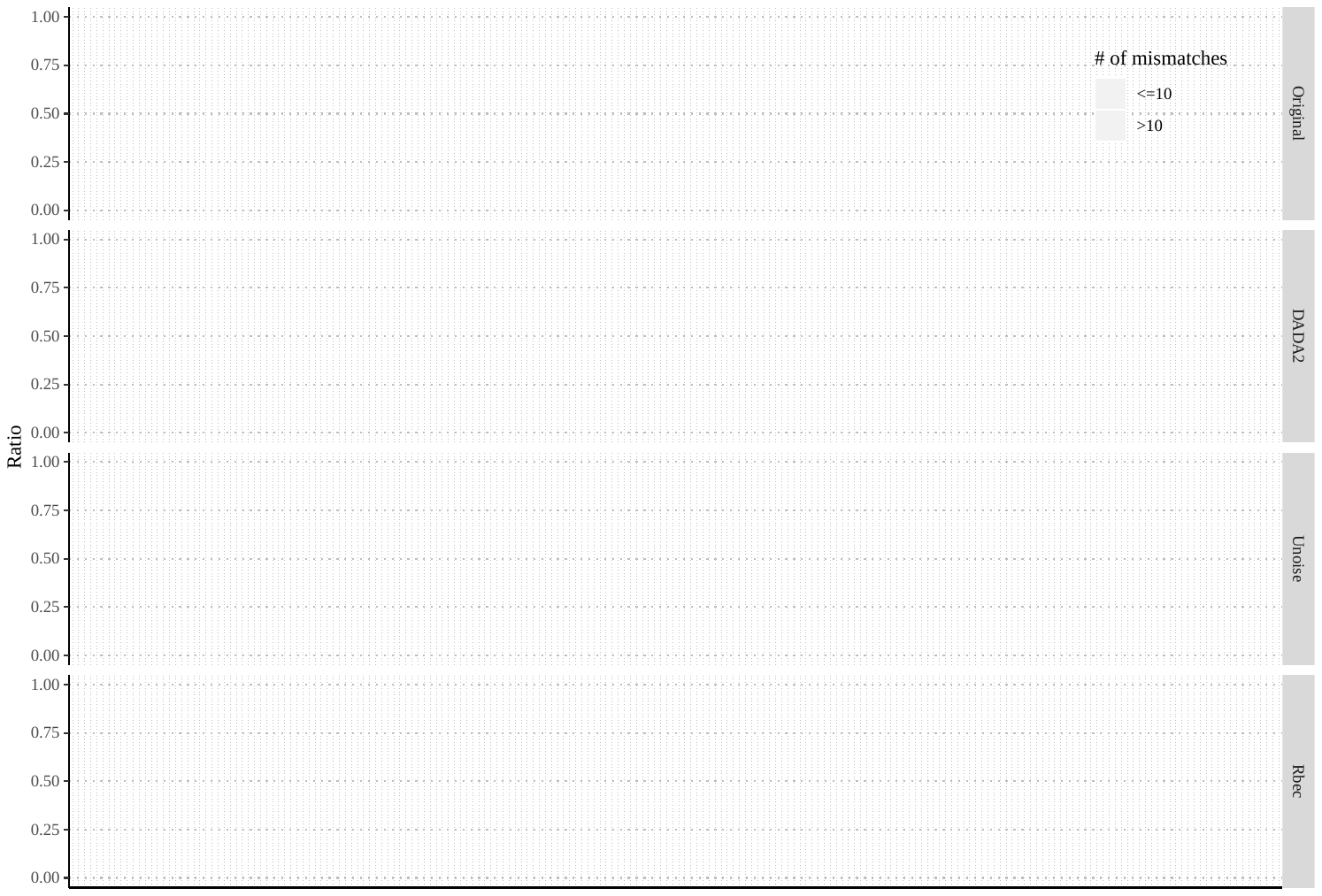

**Fig S3. Abundance ratios between second-most abundant and most abundant unique tags in each sample before and after correction with different methods.** The *x* axis represents different strains and *y* axis represents the abundance ratio calculated as $\frac{Abundance of second-most abundant unique tag}{Abundance of most abundant unique tag}$. The colour of each dot indicates the sequence dissimilarity between the two unique tags. Data points are depicted in dark brown if the two tags are close (≤ 10 base mismatches).

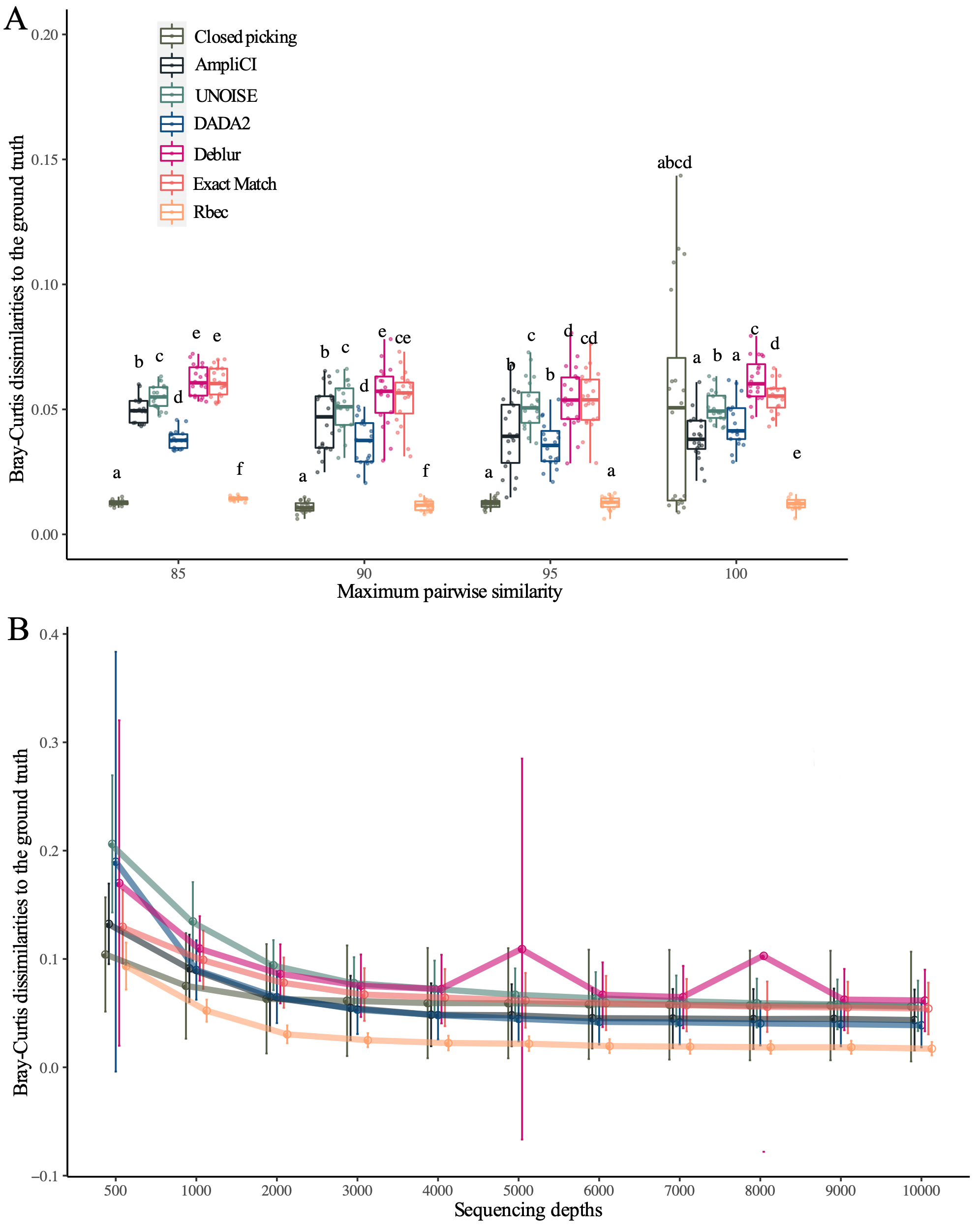

**Fig S4. Effect of strain relatedness (A) and sequencing depth (B) on the performance of different methods to characterize bacterial communities.** (**A**) Mock samples were generated from 20 strains with different strain similarities and 20 replicates for each threshold. Letters indicate the significant groups (paired Wilcox test, *P* < 0.05) within mocks with the same strain similarity. (**B**) The mock samples with different sequencing depth were directly subsampled from the mock samples with 50 strains. Circles represent the means of each method and parameter combination, while vertical lines represent the standard deviation.

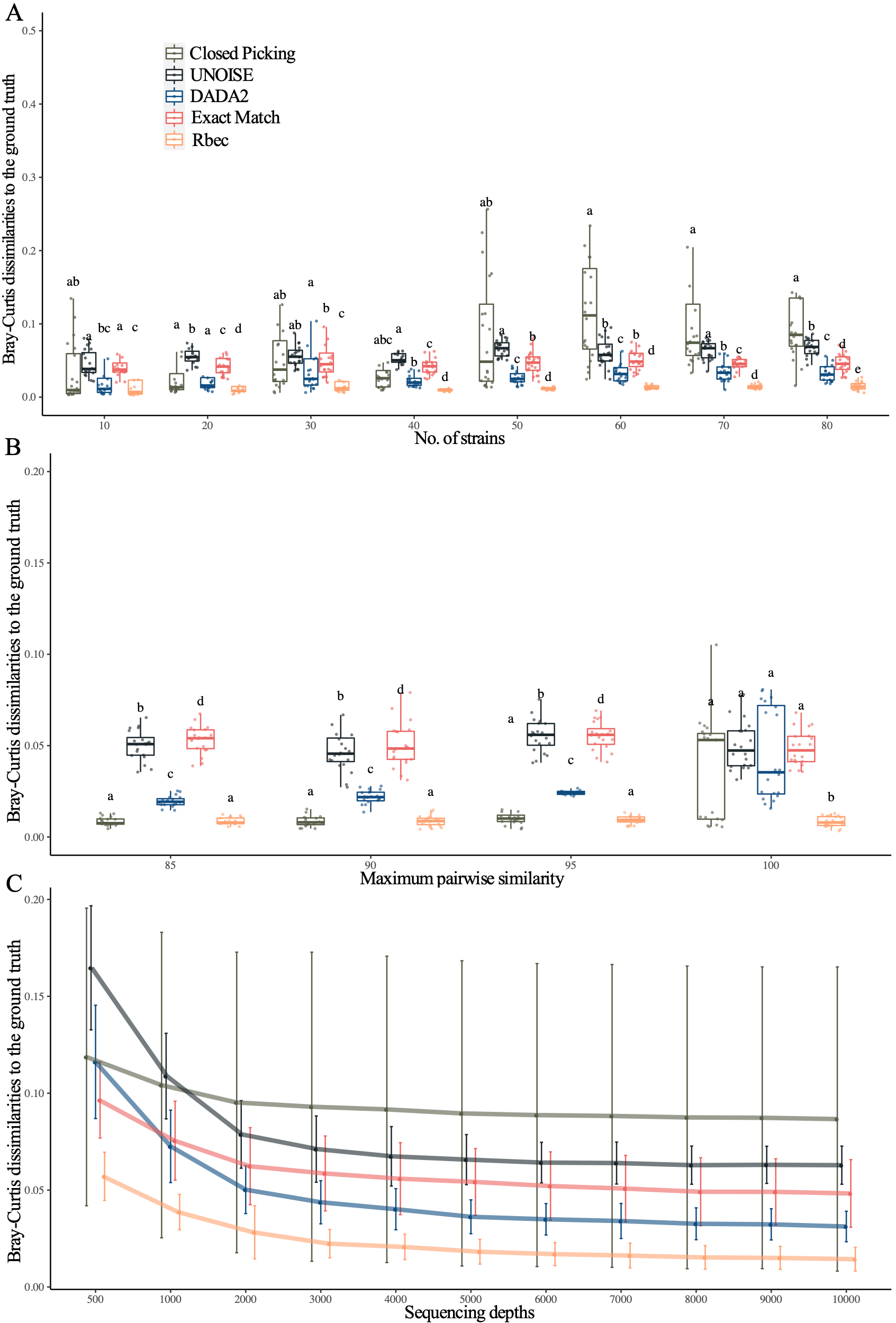

**Fig S5. Effect of community complexity (A), strain relatedness (B) and sequencing depth (C) on the performance of different methods to characterize fungal communities.** Mock samples were generated by randomly picking fungal strains from 97 candidates and mixing subsampled reads from the corresponding strains. For mock samples with different similarities, only 20 fungal strains were included for each mock, while mock samples with different sequencing depths were directly subsampled from the dataset containing 50 fungal strains. For each threshold, 20 replicates were generated. Letters indicate the significant groups (paired Wilcox test, *P* < 0.05) within mock samples with the same parameters.

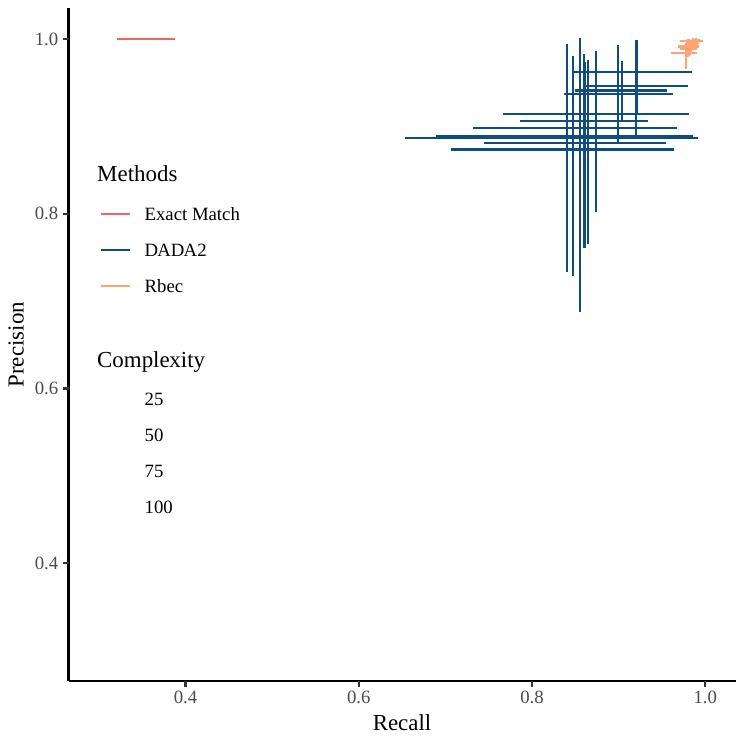

**Fig S6. Precision and recall of different methods.** Bacterial mock communities with different complexities were analysed to calculate the precision and recall of different methods. We used the formulas $Recall=\frac{No. of precisely corrected \mathrm{reads}}{Total No. of reads}$ and $Precision=\frac{No. of precisely corrected reads}{No. of corrected reads}$. Crossed vertical and horizontal lines represent the standard deviation of precision and recall in each complexity threshold, while dots indicate the mean values.

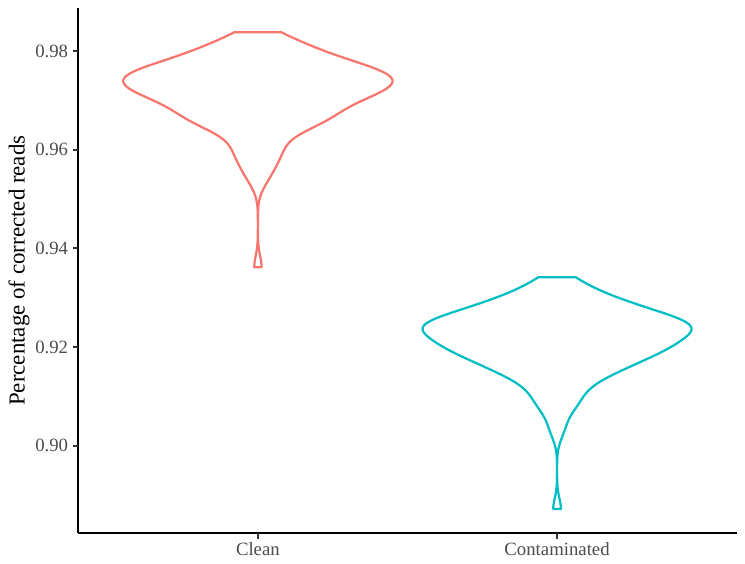

**Fig. S7. Segregation of percentages of corrected reads between clean and contaminated SynCom samples.** A set of 100 ‘clean’ mock communities were generated by randomly picking up 50 bacteria from the bacterial seed pool and mixing the subsampled reads from corresponding strains. To generate a comparable set of contaminated samples, amplicon reads from the *E. coli* K12 *16S* rRNA sequence were added to each clean mock community to make up 5% relative abundance.
